# A Novel Set of Cas9 Fusion Proteins to stimulate Homologous Recombination: Cas9-HRs

**DOI:** 10.1101/2020.05.17.100677

**Authors:** Chris R. Hackley

## Abstract

CRISPR/Cas9 has revolutionized genetic engineering, however, the inability to control double strand break (DSB) repair has severely limited both therapeutic and academic applications. Many attempts have been made to control DSB repair choice, however particularly in the case of larger edits, none have been able to bypass the rate-limiting step of Homologous Recombination (HR): long-range 5’ end resection. Here we describe a novel set of Cas9 fusions, Cas9-HRs, designed to bypass the rate-limiting step of HR repair by simultaneously coupling initial and long-range end resection. Cas9-HRs can increase the rate HR by 2-2.5 fold and decrease cellular toxicity by 2-4 fold compared to Cas9 in mammalian cells with minimal apparent editing site bias, thus making Cas9-HRs an attractive option for applications demanding increased HDR rates for long inserts and/or reduced p53 pathway activation.

**Summary:** A novel Cas9 fusion protein designed to increase HDR rates through bypassing the rate limiting step of homologous recombination repair.

## Introduction

CRIPSR/Cas9 technology has revolutionized genetic engineering by allowing quick and fascicle targeting of virtually any accessible spot in the genome via RNA based guides^1–3^. CRIPSR/Cas9 generally has two broad uses in genome editing: mutation of targeted sites via imprecise Non-Homologous End Joining (NHEJ) mediated repair and introducing precise edits in the genome ranging from 1 to 1000s of base-pairs (bp) using an external template sequence via Homology Directed Repair (HDR)^4^. While HDR methods generally allow for more flexible editing, NHEJ repair is highly preferred in higher eukaryotes where HDR repair rates can be as much as two orders of magnitude less than NHEJ repair^5^. Low HDR/INDEL ratios have so far generally limited therapeutic applications of CRISPR/Cas9 to only those which are amenable to NHEJ based repair, and makes the introduction of targeted mutations, insertions or deletions difficult, expensive and time consuming^6,7^. In addition, NHEJ based repair can activate the p53 pathway, prolonged activation of which can lead to cellular apoptosis, not only reducing yields of edited cells but also potentially selecting for cancerous mutations^8^.

There are many different approaches that have been taken to attempt to improve HDR rates, however most fall into four broad classes: small molecule and genetically encoded inhibitors or activators of NHEJ and HDR respectively, cell cycle synchronization, optimization of culture conditions, and engineering of the nuclease(s) themselves. For example, inhibiting DNA ligase IV, a key step in the NHEJ repair pathway, or stimulating Cyclin D1, which governs the transition from G_0_ to G1/S phase, can increase HDR 2-3 fold^9,10^. While these can be effective in-vitro, these techniques must be optimized for each cell type, and can have undesirable side effects which significantly limit their usefulness. Significant effort has been put into engineering Cas9 (and other nucleases) to provide more flexible alternatives^11^, with some of the most successful being base-editors and prime-editing, both of which demonstrated significant (>20%) error free editing for both single-base (A->G and C->T) and short (∼<40bp) insertions or deletions with minimal cellular toxicity in multiple cell types^12,13^. While there are still some issues to resolve with these tools, particularly off-target effects, there is no doubt that they have dramatically impacted the landscape of genetic engineering and point to the significant benefits that engineered Cas9 fusions can bring.

Unfortunately, Cas9 fusions designed to increase the HDR rate for longer inserts (>0.3kb) have not been as successful, though progress has been made by some groups. Recently, It has been shown that fusion of various factors involved in double strand break (DSB) repair choice (CtIP, Mre11, and a truncated piece of p53 named DN1s) to Cas9 can increase the ratio of HDR/INDEL repair for longer inserts from ∼1.5-2.5 fold, depending on fusion partner^14–16^. While these results demonstrate this type of approach can succeed, each of these fusions still has significant issues. Cas9-CtIP has been shown to have both editing site and cell type bias, limiting their potential usefulness significantly^17^. Additionally, given that both CtIP and Mre11 undergo extensive post-translational regulation and have a myriad of protein:protein interactions, it is likely that these issues are related to endogenous cellular components themselves, and if so, unlikely to be improved upon^18,19^. Additionally, none of these fusions have directly been shown to reduce cellular toxicity, in fact the most effective in terms of HDR/INDEL ratio increase, Cas9-DN1s, actually increases toxicity by roughly 5-10%^16^. Equally important, their mechanisms of action fundamentally limit their effectiveness to endogenous rates of HDR repair, as none of these fusions act on what is thought to be the rate limiting step: long range 5’->3’ end resection^19^.

Eukaryotic DSB repair takes place via two main pathways: canonical and non-canonical. The first relies on binding of MRN/CtIP complex slightly downstream of the DSB, which then resect back via 3’->5’ exonuclease activity to create short (<20bp) single strand (ss) DNA 3’ ends. After additional steps, the overhanging ssDNA 3’ ends are eventually further resected via long range 5’->3’ exonucleases Exo1 or Dna2, which then commits the DSB to be repaired via HR^17^. The non-canonical pathway simply by-passes the initial 3’->5’ resection, with either Exo1 or Dna2 directly initiating and resecting the DSB 5’ ends to create long 3’ ssDNA ends, thus committing the DSB to HR repair. It is thought that the canonical repair pathway prevails due to DBS ends being “blocked” by other bound proteins^20^, however, fusion of either Exo1 or Dna2 to Cas9 should allow them preferential access to the DSB, in theory greatly increasing the chance of committing the DSBs to HR via the non-canonical repair pathway.

While both human Exo1 and Dna2 were considered as initial fusion partners, ultimately, hExo1 was chosen as due to the greater amounts of biochemical and structural data available compared to Dna2^21,22^. Full length hExo1 is a relatively large protein at 846AA in length that can be divided into roughly two regions: the N-terminal exonuclease region (1-392), and the C-terminal region that interacts with MLH2/MSH1 DNA mismatch repair proteins (393-846)^23^. While Exo1 activity and stability is also extensively post-translationally regulated by phosphorylation (similar to CtIP, Mre11 and Dna2), it has several key differences: Exo1 functions as a monomer and all but one of the putative post-translational regulatory phosphorylation sites lie in the C-terminal region, deletion of which does not impede exonuclease activity^24–26^. Finally, the *Escherichia coli (E. coli)* homologue of Exo1, which possesses 3’->5’ exonuclease activity as opposed to 5’->3’ for hExo1, was fused to Cas9 and was shown to generate much longer deletions than traditional Cas9, indicating that successful fusion of hExo1 may also be possible^27^. Therefore, various fusions of hExo1 to were designed in order to test whether forcing cells to commit to HR repair via the non-canonical pathway could both increase HDR rates and decrease cellular toxicity while minimizing editing locus and cell type bias.

## Materials and Methods

### Genomic DNA extraction and Amplification

DNA was extracted from cells using the DNA-easy mini kit (Zymogen) per manufacturer’s instructions. DNA was then amplified with either standard taq (Bioneer), Fusion (NEB), Q5 (NEB), or PrimeSTAR GXL (Takara) using standard PCR protocols. Genome integrated H2B-RT-3’ was amplified with standard taq and primers hH2B-3’-F and -R with Tm=56°C for 35 cycles; H2B-RT-5’ was amplified using PrimeSTAR GXL and primers hH2B-5’-F and -R at Tm=65°C for 35 cycles. Genome integrated PuroRT-3’ was amplified with Phusion polymerase and Puro-Int-3’-F and -R at Tm=58°C for 35 cycles; PuroRT-5’ was amplified with Phusion polymerase and Puro-Int-5’-F and -R at Tm=56°C for 35 cycles. The HBB fragment used in exonuclease activity assays was amplified form genomic DNA extracted from untransfected K562 cells using primers HBB-G-Out-F and -R with Taq polymerase, Tm=56°C for 35 cycles.

### Cell culture and transfection

Unless otherwise noted, adherent cells (A549 or H1299) were seeded in 96 well plates and grown in either 50/50 F-12/DMEM or RPMI-1640 respectively, supplemented with 5% FBS and grown to roughly 70% confluency. Cells were then transfected with either Cal-Phos as described in Chen 2012 or Lipofectamine 3000 (Thermofischer)^28^. Fresh media was exchanged the next day and cellular viability was quantified on day 2.

K562 were grown in 24 well plates with RPMI-1640 supplemented with 5% FBS until roughly 70% confluent. At that time cells were either electroporated using the mNeon system (Invitrogen) using optimized settings for K562 cells, or lipofectamine 3000 again following manufactures instructions. Cells were then returned to 24 well plates with fresh media and grown for at least two days before being used in downstream analysis.

### Cellular toxicity Quantification

20µL of 0.15 mg/mL Resazurin solution was added to cells, which were then incubated for ∼4 hours. Cellular toxicity was quantified using a Tecan SpectraFlour Plus plate reader (Tecan) using a 535/595 filter set. Raw data was then normalized to the mean of the untransfected controls.

### H2B-mNeon Knock-in HDR quantification

K562 cells were transfected with 500 ng of Cas9-HR 4,5,6,8 or Cas9 targeting hH2B-G4, plus 50 ng of hH2B-mNeon RT. After 2 days, cells attached to coverslips coated with 0.01% poly-l-lysine, and were fixed in 4% PFA (Thermofisher) for 15 minutes at RT. After fixation, cells were washed 3X in PBS, mounted in 50% glycerol and imaged on a Nikon Eclipse E600 with standard FITC filters. For quantification, random patches of cells (usually between 8-15 cells per image) were identified via Brightfield, then fluoresce images were taken with constant illumination and exposure time (100%, and 110ms respectively). After acquiring roughly 10 images per construct, ratios of positive to total cells were calculated and plotted either in absolute or normalized to Cas9 (NT), with two independent experiments each treatment.

### Cas9-HR 3 and Cas9 protein purification

pET-28b-Cas9-HR 3,4, 8 and pET-28b-Cas9 were transformed into BL21(DE3) bacteria. Single colonies were picked and grown overnight at 37°C in LB supplemented with 75 µg/mL Carbenicillin. The next day, each was diluted was 1:100 in fresh Terrific Broth media supplemented with 75 µg/mL Carbenicillin, 0.05% Glucose, 10-50 µM IPTG and grown overnight at room temperature. Purification protocols were based on a modified version of a previously published two-step Cas9 purification protocol^29^.

### Cas9-HR and Cas9 in-vitro exonuclease assay

DNA was isolated from wildtype K562 cells, and amplified using standard Taq DNA polymerase and HBB-out-F (5’-aacgatcctgagacttccaca-3’) and HBB-out-R (5’-tgcttaccaagctgtgattcc-3’), Tm=56 for 35 cycles, and purified using the Quiagen PCR cleanup kit. Cas9-HR 3,4,8 or Cas9 were combined with the amplified HBB fragment at a 10:1 molar ratio (30nM: 3nM) in 1X Cas9 reaction Buffer (50mM Tris, 100mM NaCl, 10mM MgCl_2_, 1mM DTT, pH7.9) and incubated for 1hr at 37°C, after which 1µL of Proteinase K (NEB) was added and the reaction was incubated for an additional 20 minutes at 65°C. The samples were then electrophoresed on a standard 1% TAE agarose gel stained with gel green.

## Results

### Initial Design and Characterization

Plasmid PX330 was chosen as the expression vector, as it allows for fascicle simultaneous expression of Cas9 and gRNA, diagramed in Fig 1A^1^. A fragment of Human Exo1 (1-352), which should lack any post-translational regulation, was N-terminally fused to Cas9 with linkers of different lengths and amino acids, which are diagrammed in Fig 1B, as linker composition and length can have a profound effect on fusion protein expression levels and activity^30,31^. In addition, two different versions of Cas9 were used: one with two nucleus localizing sequences (2XNLS)-Cas9, and one with the N-terminal NLS deleted, 1XNLS-Cas9, hypothesizing that the extra NLS could interfere with proper fusion of hExo1 (Fig 1B). Finally, a directly fused hExo1-Cas9 construct was developed as well, as it is possible for linkers to have a deleterious effect on fusion protein performance. Here after the hExo1(1-352)-Cas9 fusions will be referred to Cas9-HR 1-9. After constructs were cloned and sequenced, Cas9-HRs 1-8 were then tested in Human lung carcinoma A549 cells. A549 were chosen as they are both facile to grow and transfect, cost effective in terms of media and other reagents, and importantly have retained a functional p53 gene^32^. An intergenic region on Chromosome 12 was targeted, with the idea that if Cas9-HRs can shift cells from NHEJ to HR repair, p53 pathway activation and corresponding cell death should be reduced compared to unmodified Cas9.

**Figure 1:**
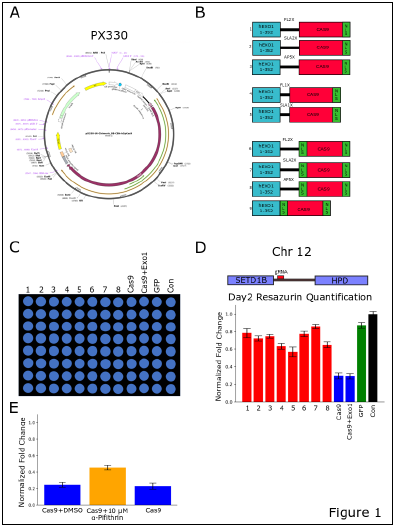
Initial Construction and Characterization of Cas9-HRs. (A) Diagram of the px330 plasmid used as the expression vector for Cas9 and Cas9-HRs. (B) Diagrams showing fusions of Cas9-HRs 1-9, with Cas9 in red, NLS sequences in green, hExo1 in teal and linkers being black lines. Sequences of linkers used are available in Table S1. (C) Example of a 96 well seeding pattern for standard A549 toxicity assays used throughout the paper. All experiments contained at least two independent replicates. (D) Cas9-HRs reduce cellular toxicity in A549 cells. The target site on Chromosome 12 is depicted above graph showing cellular toxicity for Cas9-HRs 1-8 (1-8), Cas9, Cas9+hExo1, and untransfected controls (Con). Fluorescence values were normalized to untransfected control values, two independent experiments each with 8 individual transfections. All Cas9-HRs and GFP show significant reductions in cellular toxicity relative to Cas9 or Cas9+hExo1 (p<<0.0001, two-sided students t-test). (E) α-pifithrin treatment (orange) reduces Cas9 mediated cellular toxicity compared to treatment with either solvent or Cas9 alone (both blue, p<0.001 two-sided students t-test).

Cells were plated in 96 well plates, and transfected with Cas9-HR 1-8, Cas9 and GFP using a standard Calcium Phosphate transfection technique and incubated overnight for 16-20 hours. Cells were then allowed to recover for one additional day, after which viability was cells were incubated with resazurin (MilliporeSigma) for 4 hours, after which fluorescence was quantified by a plate reader (Perkin Elmer). Figure 1D shows that all Cas9-HR fusions had greatly increased and statistically significant increased cellular viability (∼2-4 fold) compared to unmodified Cas9. Dramatically, most had similar survival rates when compared to GFP (p<<0.001 for all Cas9-HRs vs Cas9, two-sided students t-test), indicating that Cas9-HRs may in fact significantly reduce NHEJ repair and subsequent p53 pathway activation. Importantly, hExo1 fusion to Cas9 is required, as co-expression of Cas9 and hExo1 was insufficient to reduce Cas9 mediated cellular toxicity.

Next, to ensure that the Cas9 mediated toxicity was due to activation of the p53 pathway, A549 cells were again transfected with Cas9 and treated with the cell permeable p53 inhibitor α-pifithrin (MilliporeSigma). Again, after two days cellular viability was quantified via resazurin, with 10µM α-pifithrin treatment reducing cellular toxicity by ∼2-fold compared to Cas9 or Cas9 treated with DMSO (solvent). While α-pifithrin treatment could not fully reduce Cas9 toxicity, these results indicates that the cause of Cas9 toxicity in A549 cells is at least partly due to activation of p53 mediated apoptosis, the same as has been seen for other p53+ cell types^8,33^. Next, to test the assumption that toxicity is an adequate proxy for NHEJ repair rate, three new guides were designed targeting the first exon of human beta-globin (HBB, Figure S1A, right). Resazurin quantification of cellular viability of transfected A549 cells demonstrated that one guide, HBB-G3, showed significant toxicity compared to HBB-G1 or G2 (Fig S1A, left). Sanger sequencing traces of gDNA from cells transfected with HBB-G1, G2, G3 and untransfected controls demonstrated that only HBB-G3 showed the characteristic NHEJ generated INDEL pattern (Fig S1B), thus strongly indicating that cellular toxicity is an adequate proxy for NHEJ repair rate in A549 cells. Next, Cas9-HRs 1-9, Cas9 and Cas9+hExo1 targeting HBB-G3 were then transfected in A549 cells as before, with toxicity results shown in Fig S1C. Interestingly, only Cas9-HRs 5 and 9 did not show a reduction in cellular toxicity, while this time Cas9-HR 4 showed the greatest reduction. These results demonstrate the importance of modulating hExo1 positioning via linker identity and length in allowing Cas9-HRs to be functional in a wide variety of genomic loci. Given the link between toxicity, NHEJ repair rate and p53 pathways it is likely that Cas9-HRs truly reduce NHEJ repair pathway activity, however other explanations such as expression levels, mis-localization, or ablation of nuclease activity still needed to be addressed.

To rule out expression and localization issues, K562 cells in 24 well plates were transfected with Cas9-HRs 4-8 and Cas9 using lipofectamine 3000 (Thermofisher). Cas9-HRs 1-3 were omitted due to similar initial reduction in toxicity to Cas9-HRs 6-8 (Fig 1D), and Cas9-HR 9 was omitted due to lack of toxicity reduction (Fig S1C). K562 cells were used since they are p53-/-, ideally minimizing the cellular toxicity effects of Cas9 transfection and thus facilitating accurate quantification of expression levels^34^. K562 cells were either lysed with RIPA buffer (Santa Cruz) or fixed with 4% PFA, then probed for expression levels and sub-cellular localization via an α-Cas9 anti-body (Santa-Cruz) through both western blot and F-IHC. Detectable expression of full-length Cas9-HR was seen for all constructs, with generally reduced, though still detectable, expression levels for Cas9-HRs 4-7 (Fig S2A). Interestingly, Cas9-HR 8 showed similar expression levels as wildtype Cas9, again demonstrating the effects that linkers can have on protein stability and expression levels. F-IHC experiments further demonstrated detectable expression and proper nuclear localization (white arrows) of Cas9-HRs 4-8 (5-7 performed but data not shown), thus indicating that neither low, mis-expression nor improper localization are likely causes of Cas9-HRs ability to reduce cellular toxicity and NHEJ repair rates.

The next experiments were designed to simultaneously address both Cas9-HR nuclease activity and effects on HDR editing rate, as well as to further elucidate the role of p53 in Cas9 mediated cellular toxicity. First, a repair template designed to tag endongenous H2B with mNeon (referred to hereafter as H2B-mNeon RT) was constructed, including two silent mutations (indicated by the red bar) designed to disrupt Cas9 binding (Fig 2A). A549 cells were transfected using lipofectamine 3000 with Cas9-HRs 4,5,6,8 and Cas9 with H2B-G4 guides. Interestingly, Cas9-HR 8 was the only Cas9-HR to show significant reduction in cellular toxicity compared to Cas9 (Fig 2B, left p<0.001 for Cas9-HR 8, p>0.05 for all others two-sided students t-test). Next, given that α-pifithrin treatment in A549 cells likely does not fully inhibit p53 pathway activation, H1299 cells, a different lung carcinoma cell line which lacks a functional p53 gene, were used as an independent assay to further examine the effect that p53 pathway activation has on Cas9 mediated cellular toxicity. H1299 cells were plated and transfected similarly to A549 cells, and resazurin quantification of cellular viability demonstrated a dramatic reduction in toxicity for Cas9-HRs 4-6 and Cas9 in H1299 cells compared to A549, with Cas9-HR 8 only showing a small reduction (Fig 2B, right). These results demonstrate that Cas9-HR toxicity reduction is very likely due to reduced activation of the p53 pathway, indicating Cas9-HRs may be particularly useful in applications where significant p53 activation is undesirable.

**Figure 2:**
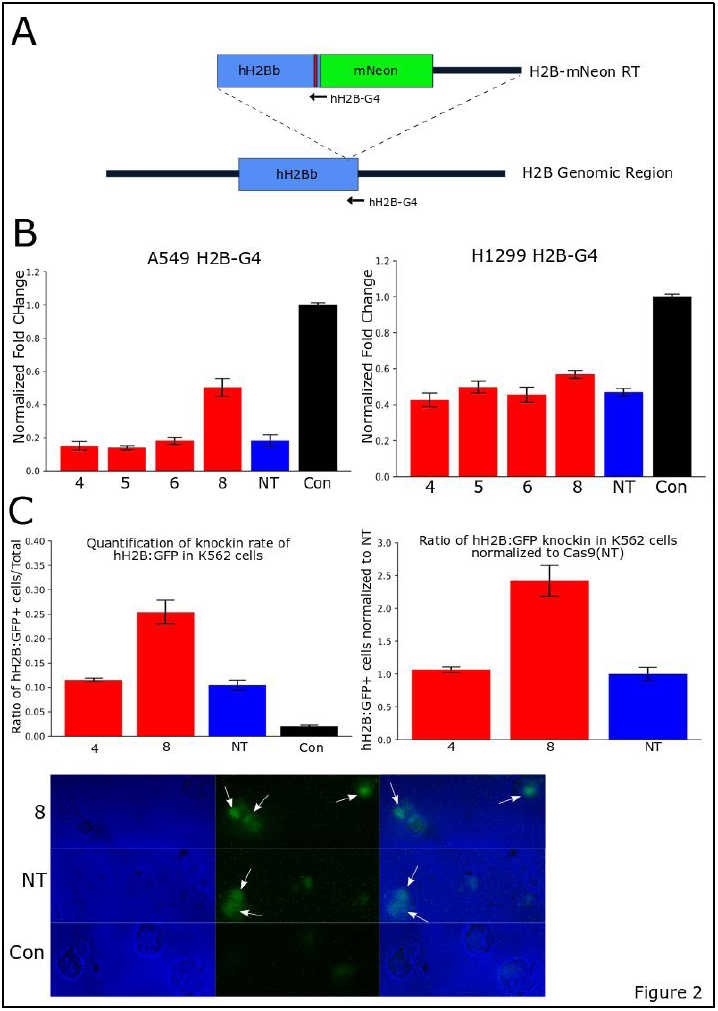
Cas9-HR 8 decreases cellular toxicity and increases HDR. (A) Diagram of the H2B-mNeon repair template, as well as showing the location of hH2B-G4 guide. Blue: H2B coding sequence, Green: mNeon, Red: silent mutations introduced in RT sequence, black lines: surrounding genomic sequence. (B) Graph of cellular toxicity of Cas9-HRs 4,5,6,8, Cas9 and untransfected controls (Con) targeting H2B-G4 in A549 cells. Only Cas9-HR shows a significant reduction in toxicity compared to Cas9 (two replicates with 8 individual transfections, p<0.001, two-sided students t-test). (C) Quantification of Cas9-HR 4,8 and Cas9 HDR rate in K562 cells. Top left and right show quantification of HDR rate; Cas9-HR 8 shows significantly higher HDR rates than either Cas9-HR 4 or Cas9 (two independent replicates, n=150-200 cells per replicate, p<0.005 two-sided students t-test). Bottom shows raw images, with Bright Field in blue (left), mNeon in green (middle), and both merged (right), with white arrows showing nuclear mNeon fluorescence.

To test whether Cas9-HRs retained nuclease activity, and if they could improve HDR repair rates, cells were transfected via electroporation with both H2B-mNeon RT and only Cas9-HRs 4,8 and Cas9, since Cas9-HRs 4 and 8 had shown the most promise in reducing toxicity. After two days, cells were fixed in 4% PFA, then directly imaged with an epifluorescent microscope, with cells showing nuclear localized fluorescence counted as successful HDR events. As with toxicity reduction in A549 cells, Cas9-HR 8 showed a significant increase in HDR rate compared to either Cas9-HR 4 or Cas9 (Figure 2C, left and right), with representative fluorescent images demonstrating HDR positive and negative cells are shown in Figure 2C, bottom. These results demonstrate that not only do Cas9-HRs retain nuclease activity, they may in fact be able to increase the HDR rate substantially.

To further confirm the image-based quantification results, DNA was extracted with the DNA-easy kit (Zymogen) from 1/10 of H2B-mNeon RT + Cas9-HR 4,8 or Cas9 transfected K562 cells. Regions surrounding the putative knock-in were amplified using 5’ (blue) and 3’ (red) specific primers (Fig S3A), with each primer pair containing one genome specific and one RT specific primer, ideally ensuring amplification of only correctly integrated HDR events. Successful amplification of both 5’ and 3’ pairs was seen with both Cas9-HR 4,8 and Cas9 (NT), but not in untransfected control samples. Additionally, the increase in band strength of Cas9-HR 8 5’ product compared to Cas9-HR 4 or Cas9 further confirms the imaging results that Cas9-HR 8 likely has an increased HDR rate relative to Cas9 (Fig S3B, 3’ appears to be out of quantitative range). The 5’ and 3’ PCR products were then gel extracted (Qiagen) and sent for sequencing (Elim Bio), where both sanger traces and consensus alignments of 4,8, and NT showed no gross differences in sequence or genomic mutations (Fig S3C). These experiments demonstrate that Cas9-HRs not only can increase the HDR rate, but as initially designed appear to maintain functionality across different cell types.

Next, an additional set of experiments were designed further test the relationship between cellular toxicity and repair pathway choice, as well as to test if Cas9-HRs can increase the rate of HDR for an independent and longer (1.8 kb) insert. First, an HDR template containing a Puromycin antibiotic resistance cassette was created via fusion PCR, with 5’ and 3’ homology arms added respectively as shown in Fig 3A, left. The target integration site is approximately ∼1kb to the 3’ end of the human H2B gene on Chromosome 6, a region which has no predicted genes or long non-coding RNA. Quantification of toxicity in A549 cells was again determined as in Fig 1. Again, only Cas9-HR 4 and 8 were tested for these experiments. Each well of A549 cells was transfected via Cal-Phos with either G-2 or G-3 targets, designed to additionally test if Cas9-HRs show any strand bias. Significant Cas9 mediated toxicity was seen with both Int G-2 and G-3, while Cas9-HRs 4 and 8 both showed a dramatic reduction in cellular toxicity (Fig 3A, right). Interestingly, Cas9-HR 4 showed significantly more toxicity with G-2 than G-3, further indicating a potential differential site preference of Cas9-HR 4 compared to Cas9-HR 8.

**Figure 3:**
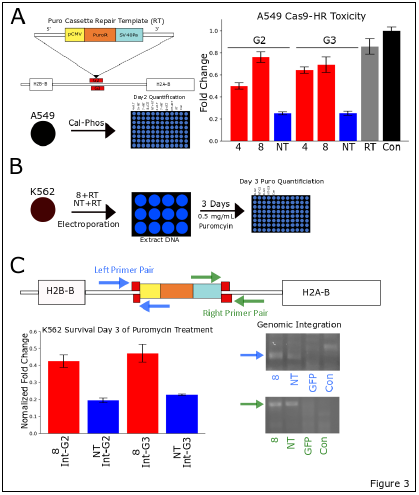
Cas9-HR 8 again shows decreased toxicity and increased HDR rates in an independent assay. (A) Left, diagram showing the Puromycin RT template, and guides Int-G2 and G3. CMV promoter: yellow, PuroR CDS: orange, SV40 polyA sequence: light blue, guide targets: red, surrounding genomic sequence: white. Right, graph showing cellular toxicity of Cas9-HRs 4,8, Cas9, PuroR RT and untransfected controls (Con) in A549 cells. Both Cas9-HR 4 and 8 show significant reductions in toxicity relative to Cas9 (two independent replicates with 8 individual transfections per replicate, p<<0.001, two-sided student t-test). (B) Experimental protocol to measure HDR rate of Cas9-HR and Cas9 in K562 cells. (C) Top, Diagram showing successful integration of PuroR RT transgene, left primer pair shown in blue, right in green. Left, bottom shows cellular viability normalized to transfection with a plasmid containing the puromycin RT. Cas9-HR 8 shows a roughly two fold increase in cellular viability compared to Cas9 with both Int-G2 or G3 (two independent replicates, 4 independent quantifications per replicate, p<0.005, student two-sided t-test. Bottom right, specific amplification is seen for both Cas9-HR 8 and Cas9, but not cells transfected with GFP or untransfected cells (Con).

K562 cells were again used instead of A549 cells, as the lack of a functional p53 gene should help to deconvolute HDR rates from cellular toxicity effects. Additionally, only Cas9-HR 8 was assayed as given previous results, it was unlikely Cas9-HR 4 would be superior to Cas9-HR 8 at either locus. K562 cells were grown in 24 well plates, which were then electroporated with Cas9-HRs 8 or Cas9 and 100ng of amplified repair template (RT), as shown in Figure 3B. After two days, DNA was extracted from ∼1/10 of surviving cells and used for analysis of Puro RT genomic integration. The next day, 0.5 mg/mL puromycin was added, and after three days cellular survival was quantified with the standard resazurin assay as before. As shown in Figure 3C (left), Cas9-HR 8 G-2 and G-3 had ∼2-fold greater surviving viable cells compared to unmodified Cas9 (NT). Again, amplification using primers designed specific for the genome (containing no sequence used in the RT) and specific for the RT at both 5’ and 3’ ends demonstrated successful integration of the repair template in both Cas9-HR 8 and Cas9, but not in either GFP or untransfected cells. These results strongly indicate that the reduction in toxicity of the Cas9-HR series of constructs is not due to lack of nuclease activity, and in-fact the Cas9-HR series likely has a significantly higher HDR editing efficiency than Cas9.

Finally, given the increasing popularity of RNP applications, protocols for bacterial expression and purification of Cas9-HRs 3,4,8 were developed. As RNP preparations of Cas9 generally lack the N-terminal NLS, Cas9-HR 3 was included due to test if the N-terminal NLS would interfere with Cas9-HR 8 purification, and Cas9-HR 4 due to potential differential site preference (Fig S1C). Successful purification of Cas9-HR 3 is shown with a representative SDS-PAGE gel in Figure 4A, with modified protocols also allowing for successful purification of Cas9-HR 4 and 8 (data not shown). Exonuclease activity assays were performed using an amplified region surrounding Exon1 of HBB (Fig 4B) as a simple first pass to test for catalytic activity of purified Cas9-HRs. Both Cas9-HR 3 and Cas9-HR 8 showed a significant reduction in the full length HBB amplicon compared to the other treatment conditions and controls (Fig 4C). These results are consistent with all plasmid based in-vivo experimental results generated so far and are a promising first step in extending Cas9 fusion protein based HDR improvement to RNP based methods.

**Figure 4:**
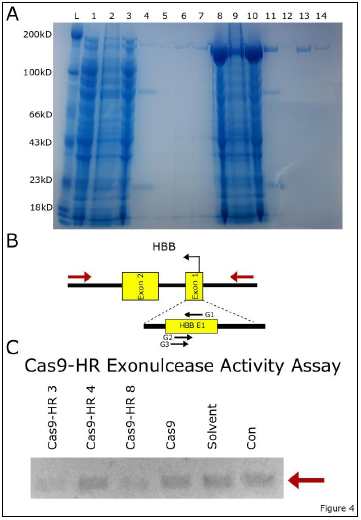
Enzymatically active Cas9-HRs can be purified from *E. Coli*. (A) SDS-PAGE gel showing successful purification of Cas9-HR 3. Lanes 1-7 Cas9-HR 3, Lanes 8-15 Cas9. Lanes 1,8 whole lysate; 2,9 soluble fraction; 3,10 insoluble fraction; 4,11 His-purification; 5,12 Sepharose wash; 6,13 Sepharose Elution 1; 7,14 Sepharose Elution 2. (B) Diagram of HBB genomic region, with primers in red showing the amplified fragment (956bp). (C) Agarose Gel image of exonuclease activity of purified Cas9-HRs 3,4,8, Cas9, Solvent (Cas9-HR buffer), and DNA (Con). Reduction in band intensity of Cas9-HR 3 and 8 indicates exonuclease activity.

## Discussion

The ability to control DSB repair pathway choice has been long sought after, with numerous different tools and techniques developed to influence repair choice^4,5,9^. The data shown here demonstrates that the first ever example of a tool successfully bypassing the rate limiting step of HR repair. Somewhat surprisingly, Cas9-HR mediated committal to HR repair via the non-canonical repair pathway seemed surprisingly effective, with cellular toxicity reduced to levels seen with GFP. Unsurprisingly, target location does affect Cas9-HR performance, however a combination of Cas9-HRs 4 and 8 were able to significantly reduce toxicity at every loci tested and showed roughly equivalent activity at a given locus across cell types, indicating Cas9-HRs are functioning as intended. Given these extensive reductions in toxicity, it will also be interesting to test different types of repair templates and transfection techniques in order to test whether the HDR rates seen here are the true limits of Cas9-HR platform, or if as the toxicity data indicates, may be limited by the repair template type (PCR amplified dsDNA) used in these experiments. Additionally, bacterial expression and purification of Cas9-HRs appears to be viable, with further testing of Cas9-HRs RNP in-vivo performance to be detailed in a later publication. Extension of Cas9-HRs to RNP based methods would be a first for any HDR stimulating fusion, and based on behavior of Cas9 RNPs, could allow for significantly higher HDR rates than reported here.

In addition to increased HDR rates, Cas9-HRs also show minimal activation of the p53 pathway, potentially allowing for extension of high efficiency HDR methods to more sensitive cell types, or even select in-vivo applications. While the current size of Cas9-HRs (∼6kb) puts them slightly too large for current adenovirus techniques, fusions using minimal Cas9s or other more compact RNA guided nucleases will be investigated to make Cas9-HRs compatible with viral techniques. Nevertheless, even in their current state the Cas9-HR platform represents a significant step forward in controlling DSB repair choice and should prove particularly useful for applications demanding increased HDR rates for long inserts and/or reduced p53 pathway activation.

