## Supplemental Data for "A Novel Set of Cas9 Fusion Proteins to stimulate Homologous Recombination: Cas9-HRs"

**Figure 1:**

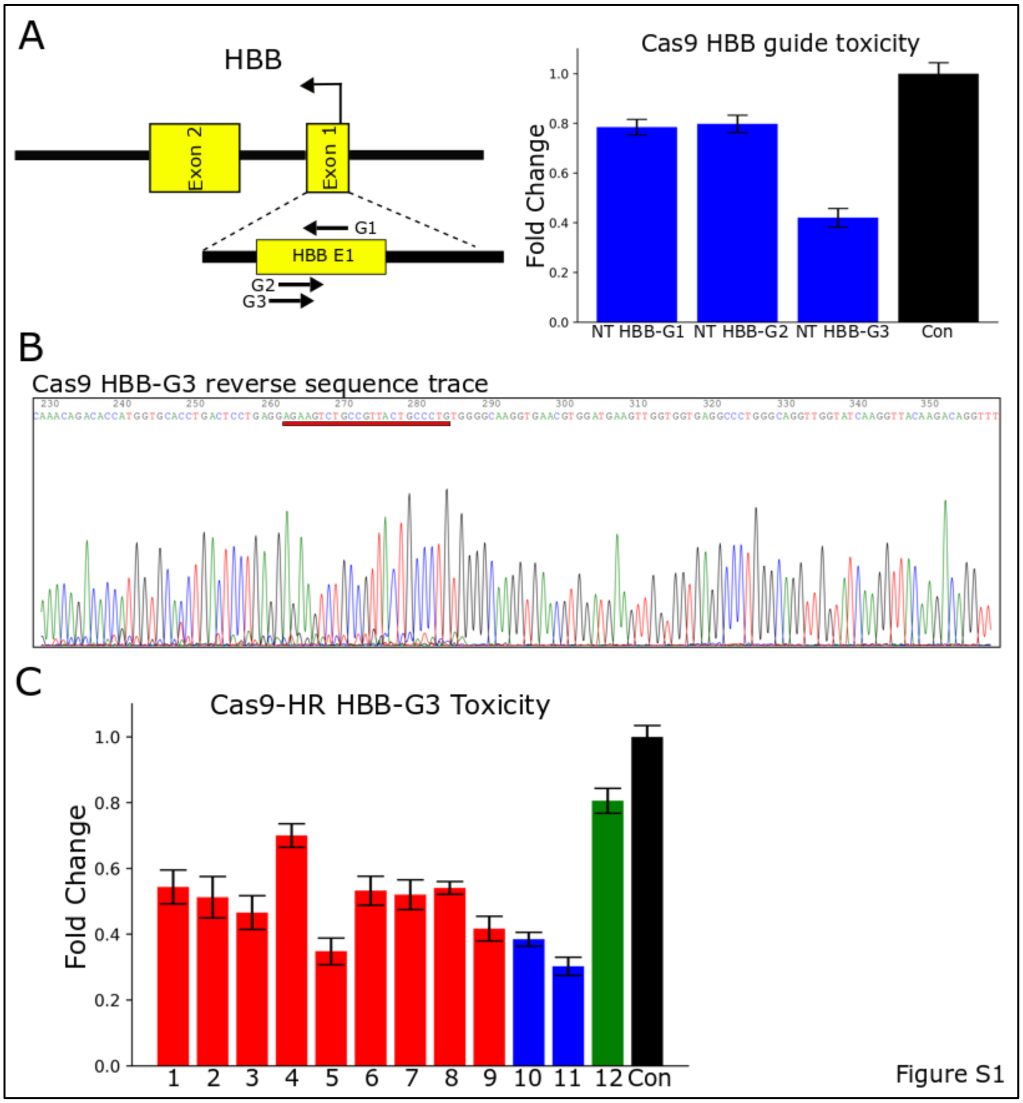


Figure S1: Cas9 mediated cellular toxicity correlates with NHEJ repair pathway activation in A549 cells. (A)Left, diagram of Human HBB exons 1, 2 and surrounding genomic sequence. Guides G1, G2, G3 shown with arrows, Exons 1 and 2: yellow, surrounding genomic sequences: black. Right, cellular viability of A549 cells transfected with Cas9 targeting either HBB- G1, G2, or G3. (B) Sequencing trace of HBB exon1 amplified from Cas9 HBB-G3 transfected cells. Red bar shows HBB-G3, showing the characteristic pattern indicating NHEJ repair. (C) Cellular viability of A549 cells transfected with Cas9-HRs 1-9 (1-9, Red), Cas9 (10, Blue), Cas9+hExo1(11, Blue), GFP (12, Green), or untransfected controls (Con, Black). Cas9-HRs 1-4, and 6-8 show significant reductions in cellular toxicity compared to Cas9 or Cas9+hExo1, $p<0.01$, two-sided students t-test.

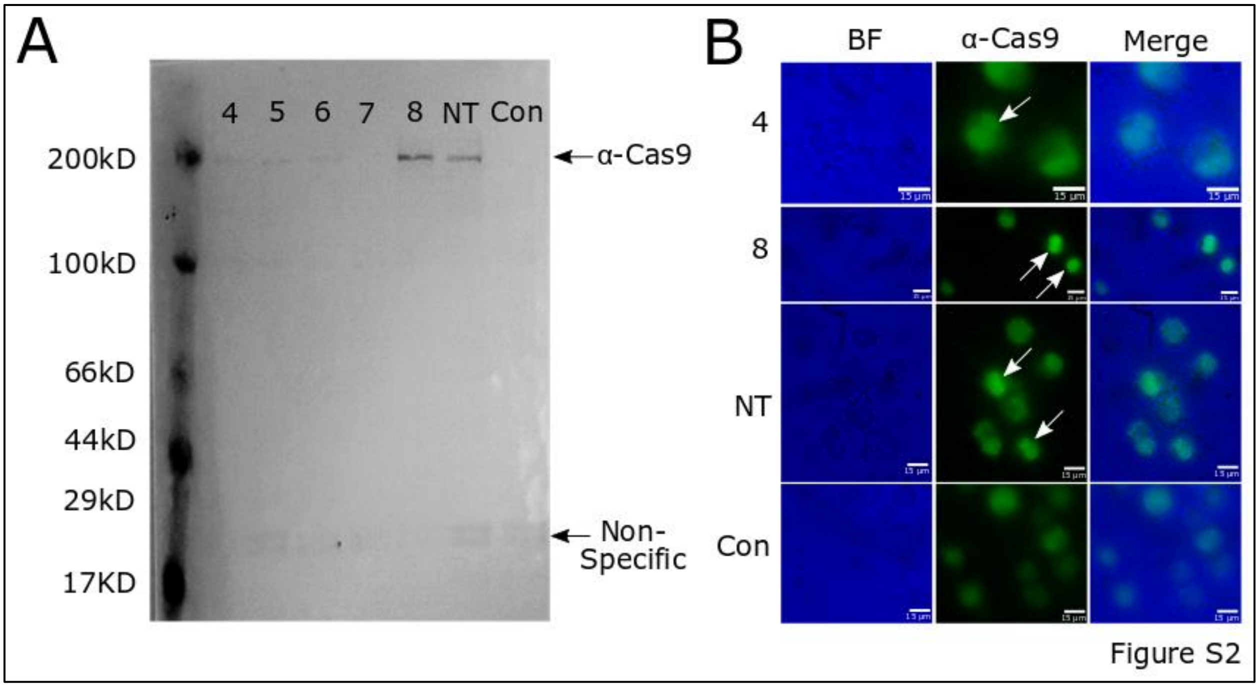


Figure S2: Cas9-HRs show similar expression and localization as Cas9. (A)Western blot from K562 cells transfected with Cas9-HRs 4-8, Cas9 (NT) or untransfected controls (Con). Black arrows show specific and nonspecific staining, as labeled. (B) Images of Cas9-HR 4, 8, Cas9 (NT) transfected or untransfected control (Con) K562 cells stained for Cas9 expression. Strong localization of Cas9-HRs 4,8 and Cas9 can be seen in the nucleus (white arrows), whereas control cells only show weak and diffuse signal.

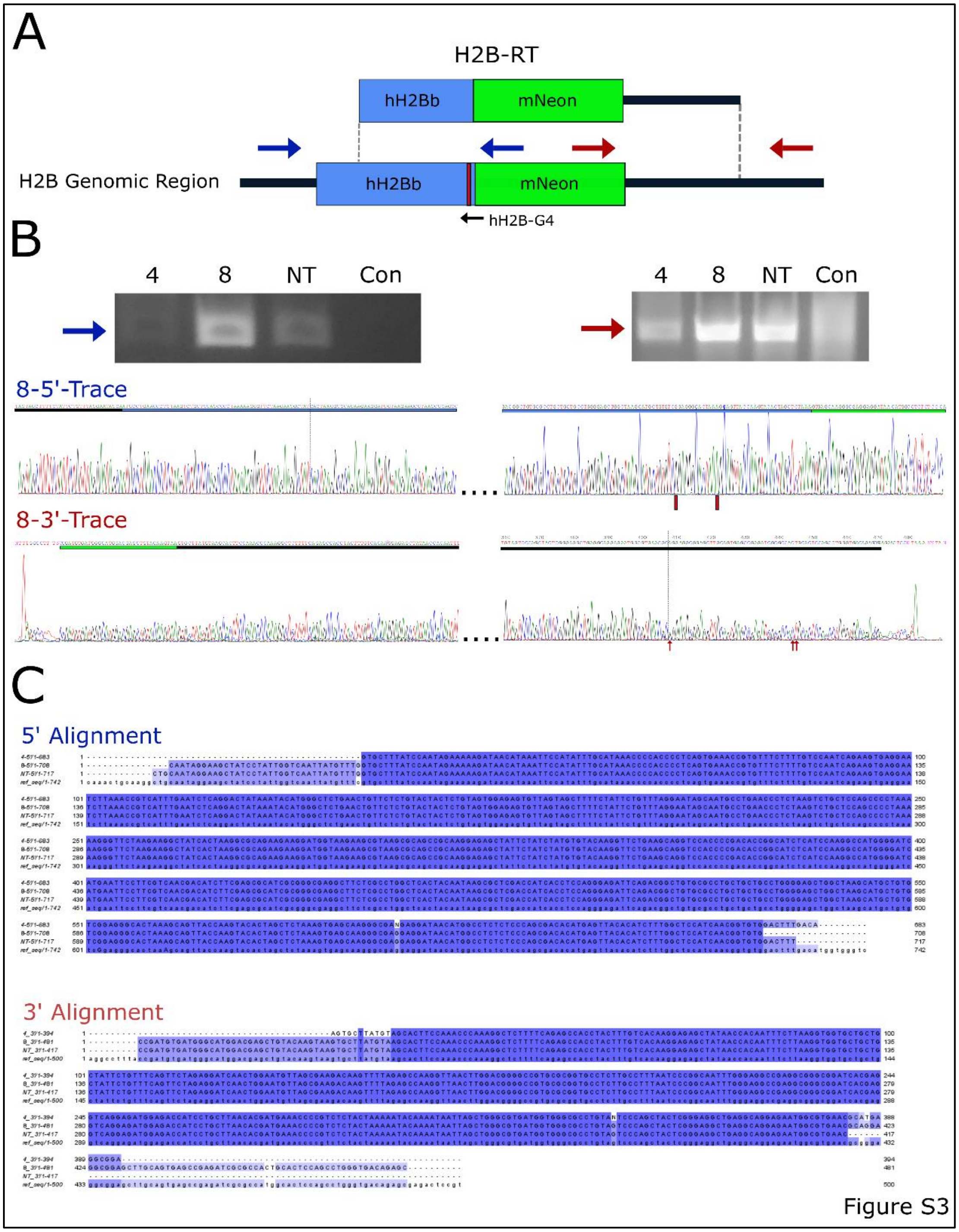


Figure S3: Genomic integration with Cas9-HR shows no detectable biases compared to Cas9. (A) Diagram showing the H2B-mNeon RT; 5' primers: blue, 3' primers: red. (B) Top left, right: Agarose gel images showing successful amplification of 5' and 3' PCR productions from gDNA isolated from K562 cells transfected with Cas9-HR 4, 8, Cas9 and H2B-mNeon RT, but not from untransfected control cells (Con). Middle, bottom sanger sequencing traces from gel purified 5' and 3' products from Cas9-HR 8. Both show no gross sequence abnormalities, Red bars :silent mutations added in the guide sequence, Red arrows: putative genomic SNPs, Black bars:

genomic sequences, Blue: hH2Bb CDS, and Green: mNeon CDS. (C) Sequence consensus alignments from sequenced PCR products form Cas9-HRs 4,8 and Cas9 when aligned to the putative integrated repair template and genomic sequences. Strong consensus is seen for all, indicating likely no gross repair differences exist between Cas9-HRs and Cas9.

**Supplementary Methods**:

**Cloning:**

pX330 was digested with AgeI, EcoRI and EcoRV, then electrophoresed and gel purified using standard procedures (Qiagen). Cas9-HR fusions 1-9 were created by amplifying hExo1 (1-352) and Cas9 from pTXB1 and pX330 respectively (see Table S2 for primers, Tm and cycles). Fragments were electrophoresed and gel purified using standard procedures, then stitched together using the following fusion PCR protocol: equimolar amounts of both fragments without primers were run for 10 cycles at a Tm of 58°C, after which outer primers (Table S2) were added and reaction continued for 20 cycles at a Tm of 62°C. Fragments were then electrophoresed, and correct sized fragments gel purified as before. Cas9-HR 1-9 purified fragments were then cloned into the pX330 backbone using infusion cloning (Takara). After colony picking and sequencing to confirm no mutations were present, Cas9-HRs 1-9 plasmids were purified using zymopure miniprep kit (Zymogen). Cas9-HRs 1-9 were then digested with BbsI, and gel purified as before. Guides (Table S1) were cloned in using standard protocols briefly consisting of denaturation and subsequent annealing, phosphorylation of 5' ends using PNK, then final cloning using T4 ligase. Colonies were picked and screened via PCR to ensure correct cloning of guides. Cas9-HRs 1-9 with various guides were then purified via ZymoPURE miniprep kits (Zymogen).

pET-28b Cas9-HRs 3,4,8 were created by synthesizing new *E.Coli* codon optimized hExo1 fragments including linkers. pET-28b Cas9 was digested with NcoI, then the backbone was gel purified as before. Both hExo1 3,4,8 and a N-terminal fragment of pET-28b were amplified using Phusion polymerase (Table S2 for primers, Tm, cycles). Fragments were then purified, then stitched together using a similar fusion PCR protocol as above: 10 cycles at Tm 62°C without primers, then 20 cycles at Tm 62°C. Correct sized fragments were then gel purified, then cloned into NcoI digested pET-28b using infusion as before.

**Western-Blot:**

Total protein was extracted from K562 cells transfected with 500ng of Cas9-HRs 4-8 or Cas9 using RIPA buffer supplemented with 2mM PMSF, 1mM Sodium Orthovandate, and 1mM protease cocktail inhibitor (Santa Cruz). After quantifying protein concentration via Bradford Assay (BioRad), 5µg total protein was run on a NuPage 4-12% Bis-Tris precast Gel (Thermofisher), then transferred at 30V for 1hr to a nitrocellulose membrane (Sigma) using the X-cell II blot module (Invitrogen). The membrane was the washed 2-4X for 5 minutes with PBST, blocked for 30 minutes with 5%non-fat milk, washed 2X with PBST, then incubated with α-Cas9 (1:1000, Santa Cruz), for 1 hr at room temperature, then overnight at 4°C. After washing 4-6X for minutes with PBST, the membrane was incubated with α-mouse-HRP(1:1000, Santa Cruz) for 1 hr at RT, and overnight at 4°C. After washing 2-4X for 5 minutes with PBST, the membrane was incubated with a 1X NC/DAB (thermofisher) solution for 15-30 minutes after which the gel was imaged.

**F-IHC:**

K562 cells transfected with 500 ng of either Cas9-HR 4-8 or Cas9, were attached to cloverslips coated with 0.01% Poly-l-lysine and fixed with 4%PFA (Thermofisher) for 15 minutes at RT and washed 4X for 5 minutes with PBST. Cells were blocked with 5% BSA for 30 minutes, then incubated with α-Cas9 (1:1000, Santa Cruz) for 1 hr at RT, then 4°C overnight. The next day cells were washed 4X with PBST for 5 minutes per wash, then incubated with m-IgG$_k$ BP-CFL 488 (1:1000, Santa Cruz) for 1 hour at RT, then overnight at 4°C. After an additional 4X PBST washes of 5 minutes per wash, cells were mounted in 50% glycerol, then imaged using a Nikon Eclipse E600 and standard FITC filters.

**Sequence analysis:**

A Plasmid Editor (APE, Wayne Davis) was used for all sequence analysis and to generate images of sequence traces. Alignment of sequencing results the 5' and 3' PCR product from H2B-mNeon Cas9-HRs 4,8, and Cas9 with the reference sequence. Sequences were aligned using ClustalOmega (https://www.ebi.ac.uk/Tools/msa/clustalo/) and pseudo-colored using Jalview to show percent identity of all sequences.
